# Model selection in statistical historical biogeography of Neotropical insects—the *Exophthalmus* genus complex (Curculionidae: Entiminae)

**DOI:** 10.1101/053611

**Authors:** Guanyang Zhang, Usmaan Basharat, Nicholas Matzke, Nico M. Franz

**Author notes:** Current address: The Australian National University, Canberra, Australian Capital Territory 2601, Australia (Nicholas Matzke).

## Abstract

Statistical historical biogeographical methods rely on the use of models that assume various biogeographic processes. Until recently model selection remains an explored topic and the impacts of using different models on inferring biogeographic history are poorly understood. Focusing on the Neotropical weevils in the *Exophthalmus* genus complex (Insecta: Curculionidae: Entiminae), we compare three commonly used biogeographic models – DIVA (Dispersal-Vicariance Analysis), DEC (Dispersal-Extinction-Cladogenesis) and BayArea (Bayesian Analysis of Biogeography), and examine the impact of modeling founder-event jump dispersal on biogeographic history estimation. We also investigate the biogeographic events that have shaped patterns of distributions, diversification, and endemism in this group of weevils. We sample representatives of 65 species of the *Exophthalmus* genus complex and 26 outgroup terminals from the Neotropics including Caribbean islands and mainland. We reconstruct a molecular phylogeny based on six genes and performed molecular dating using a relaxed clock with three fossil calibration points. We conduct biogeographic history estimations and compare alternative biogeographic models with the R package BioGeoBEARS. Model selection strongly favors biogeographic models that include founder-event jump dispersal. Without modeling jump dispersal, estimations based on the three biogeographic models are dramatically different, especially at early diverging nodes. When jump dispersal is modeled, however, the three biogeographic models perform similarly. Accordingly, we show that the Neotropical mainland was colonized by Caribbean species in the early Miocene, and that *in situ* diversification accounts for a majority (~75%) of the biogeographic events in the *Exophthalmus* genus complex. Our study highlights the need for testing for wide-ranging historical biogeographic processes in the study of Caribbean biogeography and the importance of comparing and selecting the best-fitting model in statistical biogeographic inferences. We demonstrate that modeling founder-event jump dispersal significantly improves the fit of the biogeographic history estimation of Caribbean and Neotropical mainland weevils. We establish that *in situ* diversification acts as a dominant biogeographic force in the evolution of the *Exophthalmus* genus complex. The colonization of the Neotropical mainland from Caribbean islands reinforces the notion that islands can be an important source of continental diversity.

## 1. INTRODUCTION

In this study we analyze the biogeography history of a large group of Neotropical weevils in the *Exophthalmus* genus complex (Insecta: Curculionidae: Entiminae), using a comparative, model based approach. In doing so, we are specifically interested in exploring and documenting the impacts of using different models on biogeographic inference, which include a particular focus on founder-event jump dispersal (FEJD) (Matzke 2013a, 2014). Methods of historical biogeographic estimations are founded upon models of geographic range evolution, and thus entail specific assumptions about biogeographic processes or “range inheritance scenarios” (Matzke, 2013a; Ree et al., 2005). These processes can be anagenetic or cladogenetic, and specify range-evolution scenarios such as dispersal (range expansion), extinction (range contraction), sympatry, vicariance, and FEJD (Matzke, 2013a; Ree & Smith, 2008). Three widely used biogeographic models, DEC (Dispersal-Extinction-Cladogenesis) (Ree & Smith, 2008; Ree et al., 2005), DIVA (Dispersal-Vicariance Analysis) (Nylander et al., 2008; Ronquist, 1997; Yu et al., 2010) and BayArea (Bayesian Analysis of Biogeography) (Landis et al., 2013), each assume a different set of biogeographic processes (see Matzke, 2013a). Until recently, previous model-based historical biogeographic studies did not engage in model selection. Statistical testing of the fit of alternative biogeographic models to the data (phylogeny and distribution) remains a developing area in historical biogeography. This forms a sharp contrast to phylogenetic studies of molecular and phenotypic evolution where model selection is a longstanding research branch with diverse solutions (Goldman, 1993; Hansen, 1997; O’Meara, 2012; Posada & Crandall, 1998). Therefore, it remains poorly understood whether the choice of model has any impact on biogeographic inferences and how to select among alternative models. Here we focus on a large insect radiation in the Caribbean and the Neotropical mainland and specifically address the issue of model selection in historical biogeographic inference. Using the R package BioGeoBEARS (Matzke, 2013b, 2014), we compare and statistically test the fit of three models - DEC, DIVA and BayArea - with regards to a new dated phylogeny and biogeographic patterns displayed by members of this lineage (the last two as DIVALIKE and BAYAREALIKE as they are likelihood implementations of the original models). We focus in particular on the significance of adding FEJD into these models, a biogeographic process that is explicit in the new modeling approaches and which may be an important driver of the diversification of our focal lineage.

Long-range dispersal is becoming widely recognized as an important mechanism of shaping organismal distributions and there is currently a re-emerging interest in this topic (e.g., Gillespie et al., 2012; de Queiroz, 2005; Schönhofer et al., 2013; Toussaint et al. 2016). One particular process of long-range dispersal is founder-event speciation (Carson, 1983; Gillespie & Roderick, 2002; Paulay & Meyer, 2002). Matzke (2014) pioneered the inclusion of FEJD into the widely used DEC model (Ree & Smith, 2008; Ree et al., 2005), thereby demonstrating the positive effects of modeling FEJD on estimations of ancestral ranges in island lineages. The “DEC+J” model (Matzke 2014) introduces the free parameter *j* (jump) for FEJD, where at the time of cladogenesis one daughter lineage inherits the ancestral range and the other “jumps” to a new area via founder-event speciation (e.g., A → A, C; or AB → AB, C). We purposefully restrict the application of the term founder-event speciation to this particular biogeographic range-inheritance scenario as specified in Matzke (2013a). We thereby wish to avoid confusing this term with the genetic mechanisms suggested by the founder-effect speciation theory. DEC+J, along with likelihood versions of DIVA and BayArea (DIVALIKE and BAYAREALIKE) are implemented in the R package BioGeoBEARS (Matzke, 2013b).

Members of the EGC are broad-nosed weevils in the subfamily Entiminae and the tribe Eustylini (Alonso-Zarazaga & Lyal, 1999). The largest genus in this complex is *Exophthalmus* Schoenherr, comprising some 95 species (Morrone, 1999; Peck, 2006). In accordance with Franz (2012), our primary taxonomic antecedent for this study, several other genera, i.e., *Chauliopleurus* Champion, *Compsoricus* Franz, *Diaprepes* Schoenherr, *Pachnaeus* Schoenherr, *Rhinospathe* Chevrolat, *Tetrabothynus* Labram & Imhoff, and *Tropirhinus* Schoenherr, are also entailed in this complex. As such the complex contains some 140 described species. Further taxonomic revision of the EGC is urgently needed but is not our present focus. Species of the EGC are relatively large-sized (5-32 mm in length) and often spectacularly colored weevils (Fig. 1). As many other entimines, the adults are foliage feeders whereas the larvae live in soil and feed on roots.

**Figure 1.**
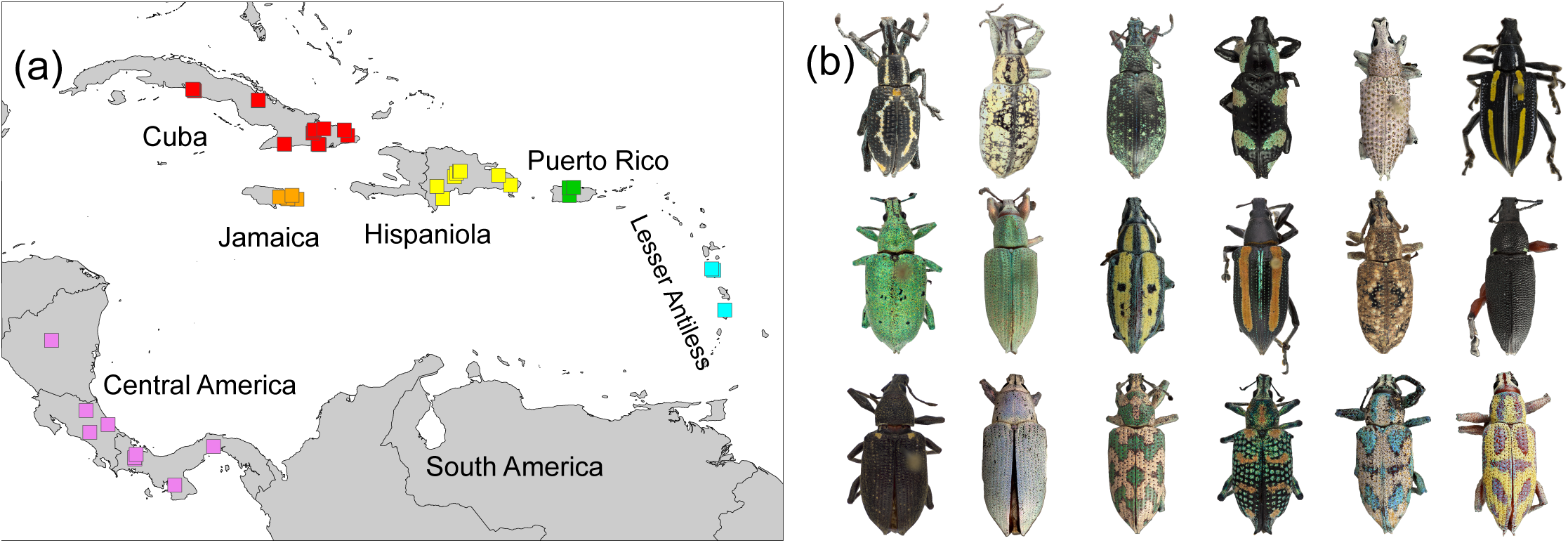
Sampling localities of the *Exophthalmus* genus complex (a) and dorsal habitus images of representative species (b) (Top: *Exophthalmus* aff. annulonotatus sp.GZ73, *Exophthalmus impositus, Exophthalmus jekelianus, Exophthalmus nr. humeridens sp.GZ49, Exophthalmus nr. viridis sp.GZ45, Exophthalmus quadrivittatus. Middle: Exophthalmus quindecimpunctatus Exophthalmus roseipes, Exophthalmus scalaris. Exophthalmus similis, Exophthalmus triangulifer, Chauliopleurus adipatus. Bottom: Diaprepes boxi, Pachnaeus sp.GZ1, Tetrabothynus spectabilis, Tropirhinus sp. 38, Tropirhinus sp.GZ46, Tropirhinus* sp.GZ47.)

Species of the EGC are distributed in both the Caribbean region and the Neotropical mainland. They are especially diverse in the Caribbean, with ~85 species or >60% of the species diversity restricted to that region. The continental species are concentrated in Central America and southern Mexico (~44 species). Only 12 species or <9% are known from South America (six exclusive to that region). The Caribbean species exhibit a high level of single-island endemism. For instance, in Cuba 19/20 recorded species of the genus *Exophthalmus* are endemic to that island (Peck, 2006), and 15/17 Hispaniolan species are endemic (Morrone, 1999).

Here we infer the ancestral range evolution of EGC species based on a newly reconstructed, fossil-calibrated, dated molecular phylogeny. Our analysis includes 65 ingroup terminals and is based on six gene fragments. Using the model selection function in the R package BioGeoBEARS, we perform statistical tests to compare the relative fit of three biogeographic models: DEC, DIVALIKE and BAYAREALIKE. We also assess the impact of modeling FEJD by adding the jump dispersal parameter “*j*” into the aforementioned models. In addition to testing effects of models and jump dispersal, we also explore answers to the following questions. What are the relative contributions of inter-island dispersal and within-island or *in situ* diversification in shaping species distributions? How did EGC species come to occupy both continental and island areas? How does the biogeographic estimation of EGC inform Caribbean biogeography?

## 2. MATERIALS AND METHODS

### 2.1 Taxonomic sampling and databasing

The ingroup taxon samples (Supplementary Table S1) include 65 species of the EGC, representing nine genera as recognized in Alonso-Zarazaga & Lyal (1999) and Franz (2012). The taxonomically widely sampled outgroup taxa include 25 species, covering several closely allied tribes: Geonemini, Naupactini, Polydrosini, and Tanymecini. Ingroup taxa were sampled from all major Greater Antillean islands; i.e. Cuba (U), Jamaica (J), Hispaniola (H), and Puerto Rico (P) - 44 species; the Lesser Antillean (L) islands of Dominica and St. Lucia-six species; and three Central American (C) countries (Panama, Costa Rica, and Nicaragua) −15 species. Sampling localities for specimens belonging to the EGC are displayed in Figure 1. The current sampling lacks specimens directly collected from South America. Based on a published morphological phylogeny (Franz, 2012) and our ongoing research, we are able to place at least seven South American species in the Central American clade of EGC (bottommost clade in Figure 2). We provide a detail analysis in supplementary method descriptions regarding the placement of South American species. These species are, however, not included in the phylogenetic or biogeographic analyses, due to a lack of molecular data.

**Figure 2.**
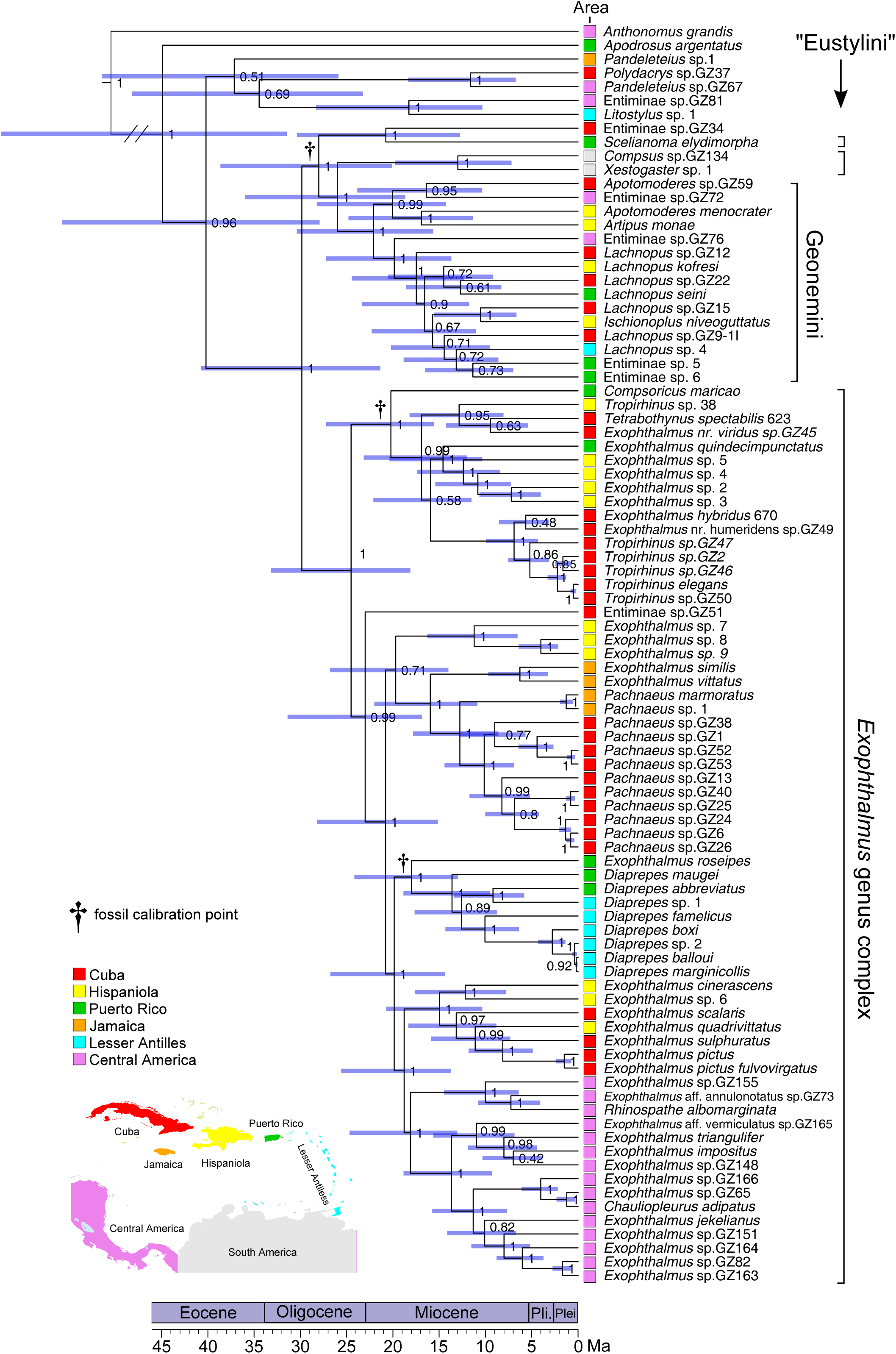
Maximum clade credibility phylogeny of the *Exophthalmus* genus complex derived from a Bayesian analysis employing a relaxed molecular clock Numbers next to nodes are posterior probabilities and horizonal bars on nodes indicate 95% intervals of ages

We identified specimens by consulting original descriptions or reference collections, sometimes with independent verifications by other weevil taxonomic experts (CW O’brien and R. Anderson). Some specimens could not be identified to the species level due to a lack of references or because they represent undescribed species. Full, standard-compliant data of DNA-extracted specimens are openly accessible through the Symbiota Collections of Arthropods Network (SCAN:http://symbiota4.acis.ufl.edu/scan/portal/index.php; see Gries et al., 2014) by looking up catalogue numbers (e.g., ASUHIC0033679; supplementary Table S1). DNA-extracted specimens or conspecifics from identical collecting events were photographed with a Visionary Digital™ Passport II system and the images made available through SCAN.

### 2.2 Molecular methods

DNA extractions were performed using the Qiagen DNeasy^®^ Blood and Tissue Kit (Qiagen). The right hind leg was excised from a specimen and extracted. Six mitochondrial, nuclear, protein-coding or ribosomal gene fragments – *Cytochrome c Oxidase subunit I* (*COI*), *Cytochrome c Oxidase subunit II* (*COII*), *12s, Arginine Kinase* (*AK*)*, Elongation Factor 1-alpha* (*Ef 1-α*), and *28s* (D2-D4 regions) – were amplified with polymerase chain reaction using EmeraldAmp MAX PCR Master Mix in an Eppendorf vapo protect thermal cycler. Primer and PCR programming information is specified in supplementary Table S2. DNA Sequencing was performed at the Arizona State University DNA Laboratory on an Applied Biosystems 3730 capillary sequencer. DNA sequences were edited with the software Geneious R7 and uploaded to Genbank (see Genbank accession numbers in supplementary Table S3).

### 2.3 Phylogenetic reconstruction and divergence time estimation

The software program MAFFT (Katoh & Standley, 2013) was used to align DNA sequences, applying the Auto alignment strategy for protein coding genes (*COI, COII, AK* and *Ef*-*1α*) and L-INS-i for ribosomal genes (*12s* and *28s*). The aligned, individual gene data sets were concatenated using SequenceMatrix (Vaidya et al., 2011) to generate a comprehensive matrix of 91 terminals and 4747 nucleotide sites. Best data partition schemes and nucleotide substitution models were determined with PartitionFinder (version 1.1.1) (Lanfear et al., 2012), using the Bayesian Information Criterion (BIC), unlinked branch lengths, and the greedy search algorithm. Three partitions and corresponding best substitution models were inferred (see details in supplementary method descriptions).

Phylogenetic reconstruction and divergence time estimation were performed in a Bayesian framework using BEAST (version 1.8.1) (Drummond & Rambaut, 2007) via the CIPRES on-line portal (www.phylo.org) (Miller, et al. 2010). The BEAST.xml file was created with BEAUTi (version 1.8.1). The concatenated data matrix was imported into BEAUTi with the three partitions inferred by PartitionFinder. Substitution models were unlinked among partitions, and clock and tree models were linked. A Birth-Death speciation process with a starting tree obtained from an initial BEAST analysis was used to inform the tree prior. An uncorrelated lognormal relaxed clock was used because initial analyses showed that the ucld.stdev values were far greater than zero, which means a strict molecular clock model could be rejected for our data. Lognormal prior age distributions were used on three fossil-calibrated nodes, which are most appropriate for calibrations based on paleontological data (Ho, 2007). Details of the prior parameter settings are provided in supplementary method descriptions. Additional analyses were performed with clock models unlinked for the three partitions or with exponential age priors. Altogether four different analyses were performed (supplemental Table S4). Early Miocene (Burdigalian; 16-21 Ma) Dominican amber fossils loaned from the University of Kansas (KU-NHM-ENT DR-888) and the National Museum of Natural History (NMNH accession numbers: Woodruff #9768 and Woodruff #9774) were used for calibration. The phylogenetic placements of these fossils were determined by coding morphological characters for matrix of Franz (2012) and running a cladistic analysis of the expanded matrix (details are provided in supplementary material). These were crown-calibrations as the fossils could be placed to the respective clades based on synapomorphic characters. Central American species of EGC were split into two clades in initial phylogenetic analyses. A small group of three Central American species clustered with an Antillean clade, supported with a low posterior probability value of 50 (supplementary Fig. S7). This result was incongruent with Franz (2012), which recovered the monophyly of Central American species based on two morphological synapomorphies. We further conducted a maximum likelihood reconstruction using RAxML, which also recovered a monophyletic Central American clade (supplementary Fig. S8). We thus constrained the monophyly of Central American species in subsequent analyses.

The Markov chain Monte Carlo (MCMC) search was conducted for 100 million generations, sampling every 10,000 generations. Convergence and stationarity were checked with Tracer (v1.6) (tree.bio.ed.ac.uk/software/tracer/) and all parameter estimates had ESS (effective sample size) values greater than 200. The maximum clade credibility (MCC) tree was generated using the TreeAnnotator (v1.8.1), after discarding the first 25% of the posterior sample of trees as burn-in. The MCC tree was annotated with median ages and 95% highest posterior density (HPD) intervals of node ages, and visualized with FigTree (v1.4.0) (Rambaut, 2014).

### 2.4 Historical biogeographic inferences and model selection

We recognize and code seven areas corresponding to Caribbean islands or larger Neotropical mainland regions: Cuba, Jamaica, Hispaniola, Puerto Rico (including the British Virgin Islands), Lesser Antilles (herein represented by Dominica and St. Lucia), Central America, and South America. Species distributions are based on primarily our collecting data, but literature records are also consulted (Morrone, 1999; Peck, 2006). Biogeographic inferences were performed with the R package BioGeoBEARS, which implements DEC, DIVALIKE and BAYAREALIKE. “LIKE” indicates that models are likelihood implementations of the DIVA and BayArea models in regards to cladogenesis assumptions. The models have two free parameters, *d* and *e*, specifying the rate of “dispersal” (range expansion) and “extinction” (range contraction) along the internal branches of a given phylogeny. They also allow specification of a set of biogeographic processes or scenarios (Matzke, 2013a: 243). BioGeoBEARS permits explicit representation of FEJD in the aforementioned models. The additional parameter *j* is written in lower case, and specifically denotes the parameter versus the process “jump dispersal”, which is denoted as “J”. The FEJD parameter can be switched on or off. Hence BioGeoBEARS can implement six specific models: DEC, DEC+J, DIVALIKE, DIVALIKE+J, BAYAREALIKE and BAYAREALIKE+J.

To test for effects of modeling FEJD, analyses were performed with and without J. The value of *j* was not pre-determined but instead was estimated anew for each analysis. Likelihood ratio tests and the Akaike Information Criterion (AIC) were used to assess the statistical significance of data-to-model fit. When nested models differ by one parameter, a statistically significant increase is indicated by approximately two or more log-likelihood units of difference in the data likelihood. The P-value was also calculated to assess whether the null model, without jump dispersal J, can be rejected. The AIC weights reflect the relative support that the data lend to the model with or without J. Similarly, separate analyses were performed under each model configuration to evaluate the relative suitability of biogeographic processes modeled in DEC, DIVALIKE, and BAYAREALIKE.

We also assessed the effect of imposing dispersal multipliers on the dispersal processes. The first (M0) allows transitions between any areas at any times (i.e., no constraint). The second (M1) applies a dispersal multiplier to model the effect of water currents on dispersal. The prevailing water currents in the Caribbean have followed a southeast to northwest direction throughout the Cenozoic (Hedges, 2001). We therefore assigned a smaller transition rate (0.5) to dispersal events in a direction against the prevailing water currents (e.g., from Cuba east to Hispaniola), and a larger value (1) to those directionally aligned with water currents (e.g., from Puerto Rico west to Hispaniola) (supplementary Table S5).

To assess the proportions of inter-island dispersal and *in situ* diversification, we counted the types of biogeographic events inferred based on the most probable ancestral ranges at the internal nodes of the ingroup taxa. We here also make a distinction between the terms inter-island dispersal and FEJD. Inter-island dispersal refers to the biogeographic event inferred from an estimation, while FEJD refers specifically to a biogeographic process built into models. FEJD could take place between two islands, but also between an island and a continent. In biogeographic estimations using models without FEJD, it is also possible to infer inter-island dispersal events, which can only be anagenetic (Ree & Smith, 2008).

## 3. RESULTS

### 3.1 Phylogenetic relationships and divergence time estimates

In the MCC phylogeny (Fig. 2), the following genera constitute a monophyletic clade: *Compsoricus, Diaprepes, Exophthalmus, Pachnaeus, Rhinospathe, Tetrabothynus*, and *Tropirhinus*. Jointly these entities correspond to the EGC in the current sense. *Exophthalmus* in the broad sense is polyphyletic. The Eustylini, as inferred herein, are polyphyletic and comprise three clades, including the EGC and two additional lineages, i.e., *Scelianoma* and the *“Compsus* sp.GZ134 to *Xestogaster* sp.1” clade.

Within the EGC, species assemblages represented on each island are not monophyletic. For instance, species inhabiting Cuba pertain to six distinct lineages and species occurring on Hispaniola or Puerto Rico are derived from five such lineages. The four endemic Jamaican species form a paraphyletic grade relative to an exclusive Cuban clade. In the MCC phylogeny where the monophyly of Central American species is not constrained, a clade containing Cuban and Hispaniolan species is nested within an otherwise exclusively Central American radiation, rendering the latter paraphyletic (supplementary Fig. S7). This nested Caribbean group *“Exophthalmus cinerascens* to *Exophthalmus pictus fulvovirgatus”* is recovered as sister to a small clade including three Central American species; however, their relationship has a low posterior probability of 50.0%. When the monophyly of the Central American species is constrained, the same Caribbean clade is inferred as sister to an exclusively Central American clade (Fig. 2).

The age estimates based on the analysis with an uncorrelated lognormal relaxed clock, linked clocked models, and lognormal age prior distributions are described here (analysis #1 in supplementary Table S4). Additional analyses showed similar age estimates for the EGC and fossil-calibrated notes, with the median ages varying by 0.58 to 4.70 Myr and the 95% HDP intervals largely overlapping (supplementary Table S4). The inferred divergence dates indicate that the diversification of the EGC began in the late Oligocene (24.6 Ma, 95% HPD: 18.0-33.4). Two daughter clades diverged within the next 1.5-4.3 Myr during the early Miocene. The Central American clade split from its sister Caribbean lineage in the early Miocene (18.1 Ma, 95% HPD: 13.2-24.7), about 6.5 Myr after the time of origin of the EGC.

### 3.2 Biogeographic history of the EGC

The biogeographic model DIVALIKE+J, with the M1 model constraint applied, yields the highest likelihood of the data among all tested models, based on the chronogram with a monophyletic Central American clade (log likelihood: LnL = −115.39; parameter estimates: *d* = 0, *e* = 0 and *j* = 0.06) (Table 1). Reconstructions that render the group of Central America species paraphyletic have consistently lower log likelihood values, regardless of the model or constraints used, and therefore not preferred. According to the best-fitting model, the ancestor of the EGC occupied Puerto Rico in the Early Miocene (24.6 Ma, Fig. 3a). One of the clade’s two descendant lineages inherited Puerto Rico as its most probable ancestral range, whereas the other lineage colonized Cuba via FEJD.

**Fig 3.**
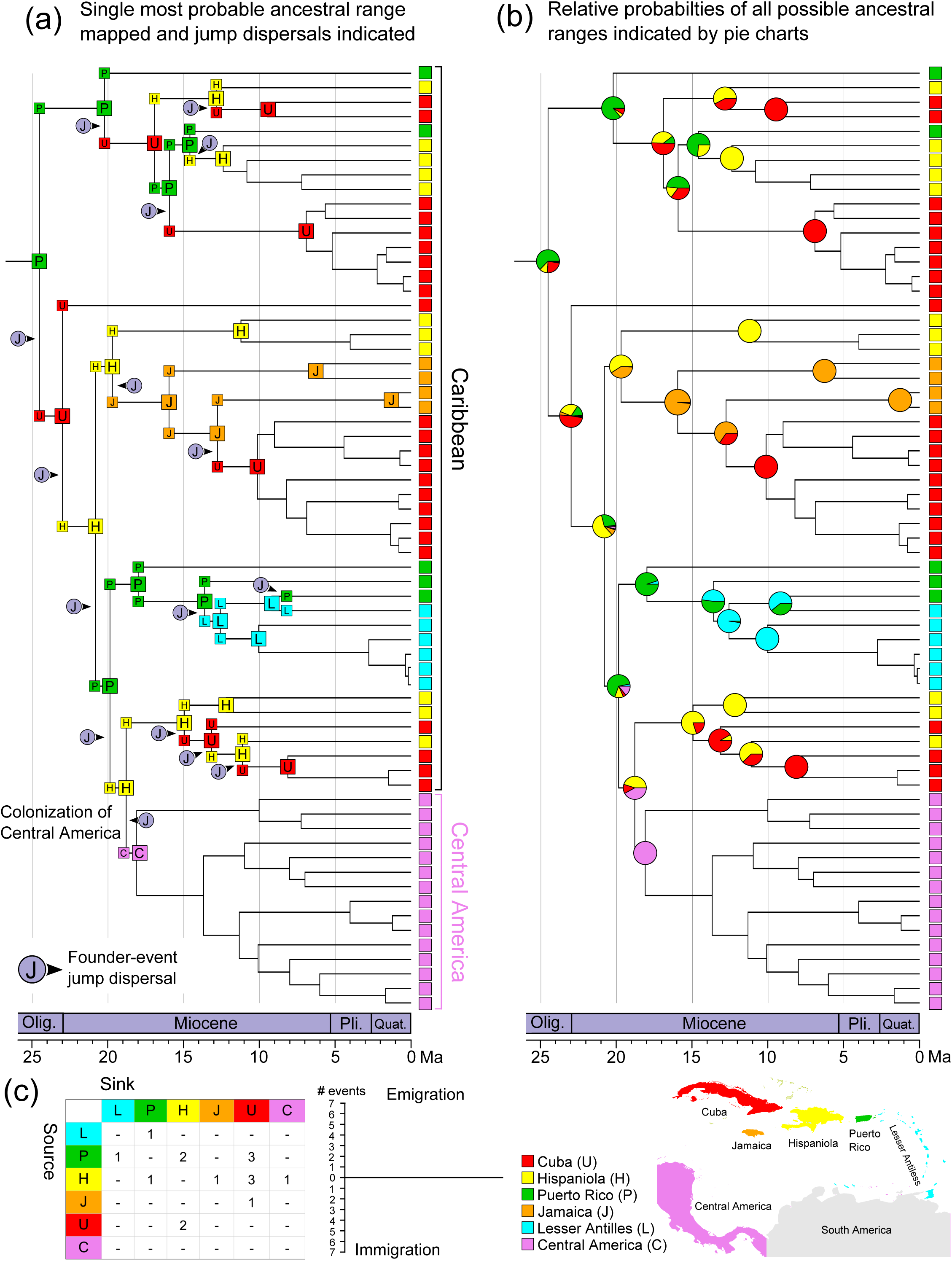
Global ancestral range estimates of the *Exophthalmus* genus complexunder the M1 constraint (i.e., dispersal rates between islands are defined according to the direction of oceanic currents) using DIVALIKE+J. The estimation was performed with BioGeoBEARS, based on the chronogram generated with BEAST. (a) Single most probable ancestral range is mapped at each node. The corner postions represent geographic range immediately after a cladogenesis event. (b) Pie charts indicate relative probabilities of all possible ancestral ranges (some of the possible ranges are not visible). Pie charts at corner positions are not shown. In both (a) and (b) the ancestral areas are omitted for nodes where no range change occured (i.e., range copying). (c) Inter-island migration direction and frequency based on most probably ancestral range estimates in (a).

**Table 1.**
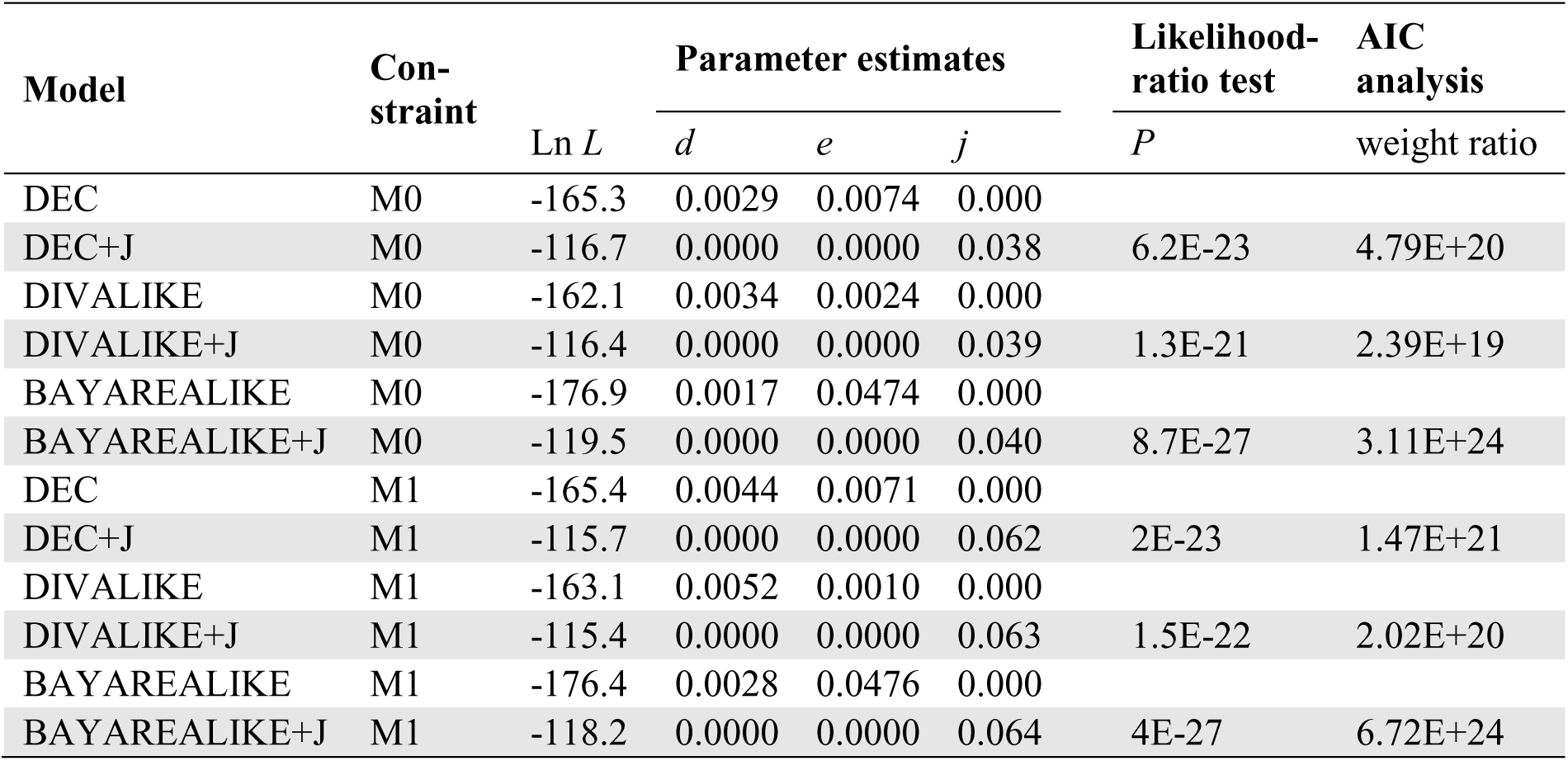
Comparison of models with and without J and two model constraints. M0 – no dispersal multiplier is applied. M1 – dispersal in the same direction of the prevailing water currents in the Caribbean is assigned a larger dispersal multiplier than that oriented against the water currents.

| Model | Con-<br>straint | Ln *L* | Parameter estimates |  |  | Likelihood-ratio test | AIC<br>analysis |
| --- | --- | --- | --- | --- | --- | --- | --- |
|  |  |  | *d* | *e* | *j* | *P* | weight ratio |
| DEC | M0 | -165.3 | 0.0029 | 0.0074 | 0.000 |  |  |
| DEC+J | M0 | -116.7 | 0.0000 | 0.0000 | 0.038 | 6.2E-23 | 4.79E+20 |
| DIVALIKE | M0 | -162.1 | 0.0034 | 0.0024 | 0.000 |  |  |
| DIVALIKE+J | M0 | -116.4 | 0.0000 | 0.0000 | 0.039 | 1.3E-21 | 2.39E+19 |
| BAYAREALIKE | M0 | -176.9 | 0.0017 | 0.0474 | 0.000 |  |  |
| BAYAREALIKE+J | M0 | -119.5 | 0.0000 | 0.0000 | 0.040 | 8.7E-27 | 3.11E+24 |
| DEC | M1 | -165.4 | 0.0044 | 0.0071 | 0.000 |  |  |
| DEC+J | M1 | -115.7 | 0.0000 | 0.0000 | 0.062 | 2E-23 | 1.47E+21 |
| DIVALIKE | M1 | -163.1 | 0.0052 | 0.0010 | 0.000 |  |  |
| DIVALIKE+J | M1 | -115.4 | 0.0000 | 0.0000 | 0.063 | 1.5E-22 | 2.02E+20 |
| BAYAREALIKE | M1 | -176.4 | 0.0028 | 0.0476 | 0.000 |  |  |
| BAYAREALIKE+J | M1 | -118.2 | 0.0000 | 0.0000 | 0.064 | 4E-27 | 6.72E+24 |

The biogeographic history of the Caribbean species is largely characterized by within-island *in situ* diversification, accompanied by relatively infrequent inter-island dispersals. A total of 16 FEJD events (25.0%) are estimated to have occurred along the 64 internal nodes in the complex, with diversification periods ranging from 24.6 to 9.2 Ma. No dispersal events occurred after 9.2 Ma (Fig. 3). *In situ* diversification is inferred for 47 of the remaining nodes (73.4%). Estimates for one node - rooting the clade *“Tropirhinus* sp.38 to *Tropirhinus* sp.GZ50” and its immediate descendants - are ambiguous. The scenario of U → H, P was plotted for that node and its two corners (Fig. 3). This scenario does not necessarily suggest a biogeographic event where the ancestral species occupying Cuba simultaneously colonized Hispaniola and Puerto Rico and disappeared from Cuba. The result merely reflects a summary of the most likely range at each of three points. The corresponding node was not counted in the calculation of biogeographic events. Colonization of Central America took place 18.8 Ma, mediated via jump dispersal from an ancestor lineage that most likely occupied Hispaniola. This event led to a subsequent radiation in Central America that would account for the origin of >40 continental species.

The reconstructed distributions, and directions and frequencies of inter-island colonization events, are shown in Fig. 3. Accordingly, Hispaniola and Puerto Rico were major source islands of colonists, accounting for 12/16 (75.0%) of out-of-island emigration events. Cuba acted as a “sink” for colonists seven times, Hispaniola four times, and Puerto Rico twice. Jamaica and the Lesser Antilles each acted once as a source or sink. Inter-island dispersal between the Lesser Antilles and the Greater Antilles was limited to Puerto Rico.

### 3.3 Model comparison and impact of FEJD

Accounting for FEJD strongly influences our analysis. Without this model parameter, biogeographic estimations based on the three biogeographic models - DEC, DIVALIKE, and BAYAREALIKE - show mixed signals. The ancestral range estimations are identical at nearly all nodes with an age less than 15 Ma (Fig. 4). These nodes involve mainly within-area diversification or sympatric range-inheritance. Between-model differences are seen mainly at deeper bifurcations. According to the DIVALIKE and BAYAREALIKE estimations, the most probable ancestral range of the EGC is “P+H+U” (Fig. 4a,c), i.e., widespread in three Greater Antillean islands. The DEC model infers “P+H+U+C”, thereby adding Central America to the ancestral range (Fig. 4b). Scenarios inferred at the intermediate nodes differ more dramatically between models. For example, in the DIVALIKE analysis, the two immediate daughter lineages connecting to the deepest node have “P” and “P+H+J+U+C” as their ancestral range, in contrast to “P+H+U” and “P+H+U+C” in DEC, and “P+H+U” and “H+U” in BAYAREALIKE.

**Figure 4.**
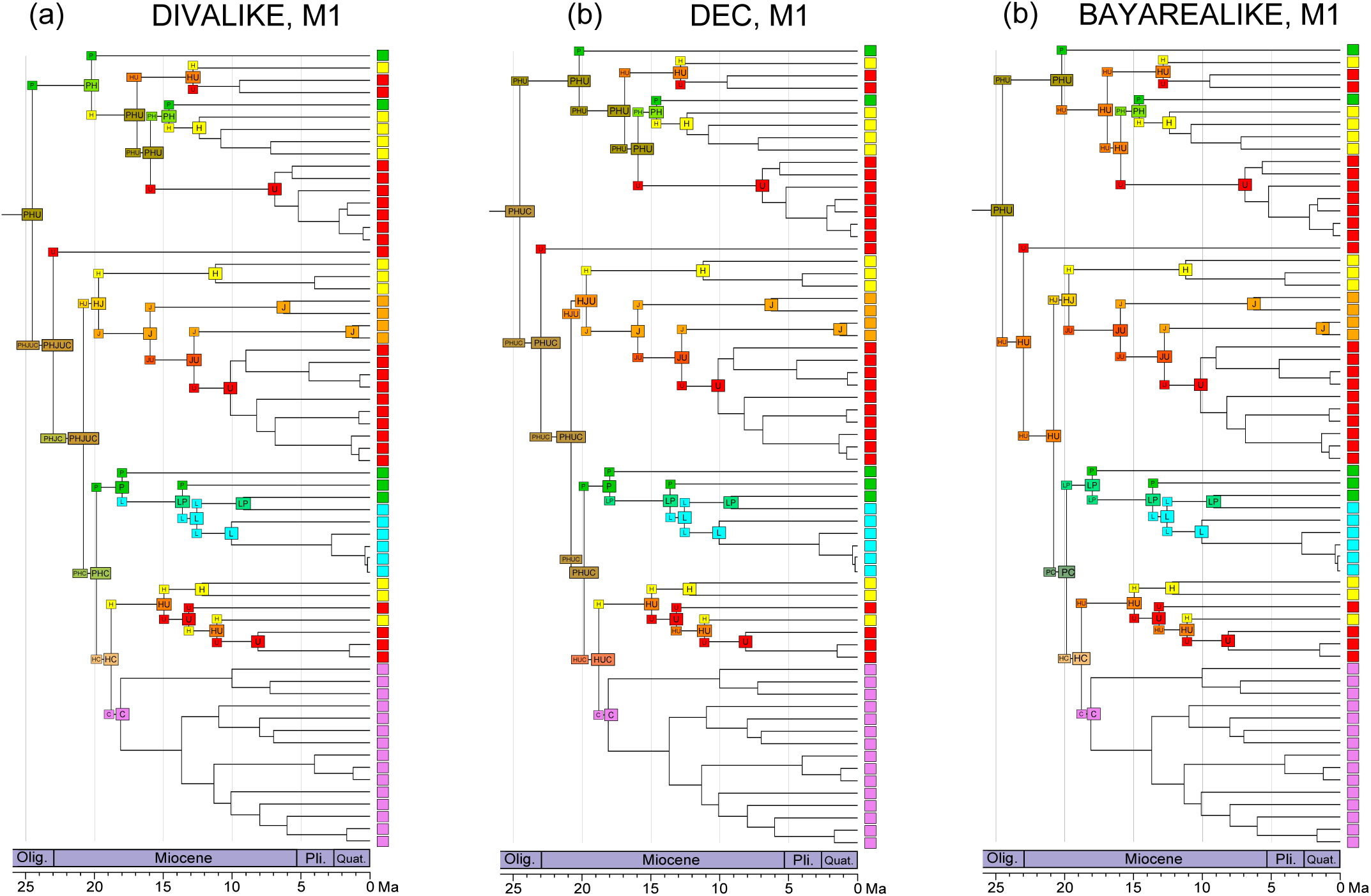
Comparison of biogeographic esitmations of the *Exophthalmus* genus complex under the M1 constraint using three biogeographic models, DIVALIKE, DEC and BAYAREALIKE, not including jump dispersal)J)h The estimations were performed with BioGeoBEARSs based on the chronogram generated with BEAST shown in Fig. 2. Single most probable ancestral range is mapped at each node. Corner positions represent geographic ranges immediately after a claogenesis event (a) DIVALIKE estimation. 4bm DEC estimation. 4cm BAYAREALIKE estimation.

Adding FEJD dramatically improves model-to-data fit. The data likelihood values obtained by adding the jump dispersal parameter *j* are 45.6–58.2 log-likelihood units higher across analytical methods or model constraints in comparison to those obtained modeling only dispersal and extinction (Table 1). The differences are statistically significant – likelihood-ratio tests for all analyses strongly favor the inclusion of *j*, whereas the null model, which states the nested model where *j* = 0 confers the same likelihood on the data, was always rejected at P < 0.0001. In addition, the AIC model weights show decisive support for models including FEJD, with exceedingly high AIC weight ratios (2.39E+19 – 6.72E+24). Accounting for FEJD also significantly affects parameter estimations: the *j* parameter is always positive, and the *d* and *e* parameters are effectively negligible. Lastly, adding FEJD to the models strongly refines the ancestral area inferences. When added, the most likely ancestral areas of all internal nodes within the ingroup are consistently resolved as single areas, whereas widespread ancestral ranges are obtained at 16 nodes in absence of this parameter (Fig. 5a,b).

**Figure 5.**
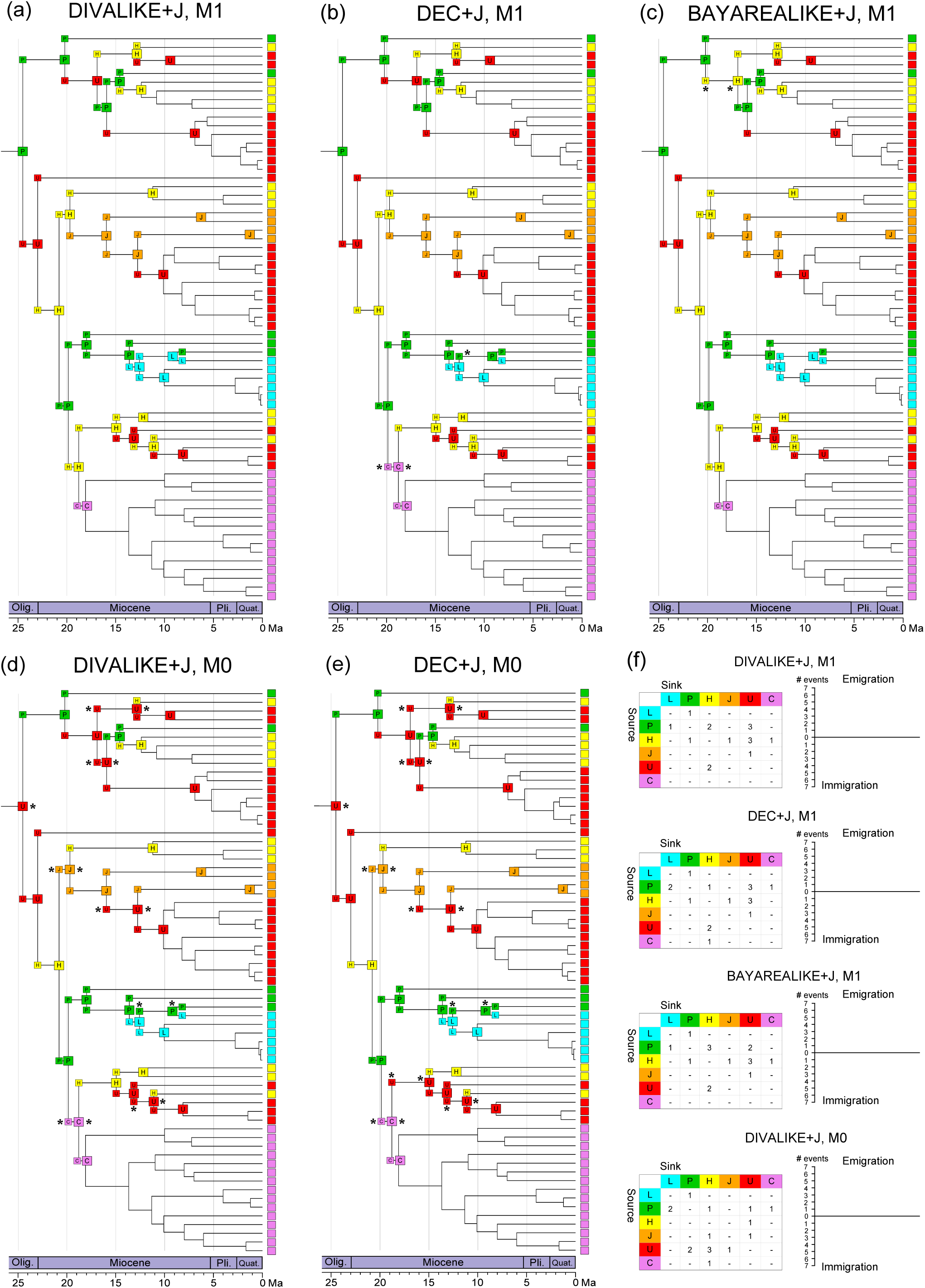
Comparison of biogeographic estimations of the *Exophthalmus* genus complex between models with and without jump dispersal, and with different model constraints (M0, M1).(a, b,c) Models that include founder-eventjump dispersal DIVALIKE+J, DEC+J and BAYAREA+J, all based on the M1 model constraint (d,e) Estimations without any specific model constraint (M0). (f) Inter-island migration directions and frequency distributions according to different models or constraints. Stars (×) next to boxes indicate different estimations from that in the “DIVALIKE+J, M1” result.

Application of the M1 model constraint and jump dispersal to the three general models yields similar biogeographic estimates (Fig. 5). Selective application of the M0 model (no constraint) versus the M1 model (favoring dispersals along prevailing water currents) primarily affects the directionality of the dispersals (Fig. 5a,c,f). Overall, these differences are too small to favor one model over another (Table 1; see also Burnham & Anderson, 2002, p. 78). When no constraint (M0) is applied, the DIVALIKE+J model estimates ten dispersals against the prevailing oceanic water currents, and six following the currents (Fig. 5f). With the M1 constraint activated for the same model, 12 dispersals are directionally aligned with water currents and four are oriented against the currents. Patterns of emigration-immigration also differ: under the M0 constraint Cuba is a major source for colonists, whereas under the M1 constraint the island acts mainly as a sink.

All biogeographic estimations under both model constraints (M0, M1), using six models (DEC, DEC+J, DIVALIKE, DIVALIKE+J, BAYAREALIKE and BAYAREALIKE+J) are shown in Figs. S1–S6 in supplementary figures.

## 4. DISCUSSION

### 4.1 Impacts of biogeographic models, FEDJ and model constraints

Our results show a complex, two-tiered pattern of data-to-model fit that demonstrates the importance of testing broadly for biogeographic models in the present and likely other Caribbean/Neotropical lineages. Most significantly, accounting for FEJD dramatically increases model fit regardless of the biogeographic method used. In biogeographic estimations with FEJD, additional non-founder dispersals and extinctions are unnecessary for explaining range evolution in the EGC. Dispersal is perfectly correlated with cladogenesis via FEJD. The three “+J” models produce highly congruent results concerning ancestral ranges and dispersal frequency and direction (Fig. 5). These results corroborate Matzke’s (2014) conclusion that biogeographic models with FEJD best account for island radiations. Our findings underscore the apparent importance of FEJD as an important mechanism shaping the biogeographic history of Caribbean weevils. This study, together with several recent historical biogeographic studies of organisms with a strong presence on oceanic islands (e.g., Berger, et al., 2016; Matos-Maraví et al., 2013; Voelker et al., 2014; Shaw et al., 2015; Tänzler et al., 2016) start to reveal an emerging pattern: FEJD provides betters model-to-data fit and acts as a critical biogeographic process for the evolution of island organisms.

The second-tier insight is that without modeling jump dispersal, the choice of biogeographic model (i.e., DEC, DIVA and BayArea) can have a major impact on ancestral range estimations, resulting in significant differences in the most probable ancestral areas and the inferred biogeographic scenarios (Fig. 4). In particular, DEC and DIVALIKE estimates yield higher data likelihoods than BAYAREALIKE (Table 1). Whereas DEC and DIVA allow for different kinds of cladogenetic events, BayArea omits all range-changing cladogenetic processes and instead assumes that all changes in geographic range occur via dispersal and extinction events along branches. However, the relatively lower data likelihoods of the BAYAREALIKE inferences indicate that cladogenesis was an important process in the biogeographic history of Caribbean weevils. In the current literature it is still commonplace to see the use of a single model for biogeographic estimation not based on explicit model selection. Since methods are now available to test and compare models, future research should be more explicit about the choice of model and its effects on biogeographic interpretations.

Assigning higher transition rates to dispersal in the same direction of the prevailing water currents in the Caribbean under the model constraint (M1) only marginally improves data likelihoods (Table 1), while allocating highly differential roles to Cuba and Hispaniola (Fig. 5a,c,f). Under the M0 model constraint, Cuba functions primarily as a source of colonists, providing emigrants to Puerto Rico, Hispaniola and Jamaica. Under that same model, Hispaniola acts as a major sink of colonists. Conversely, the M1 model constraint renders Cuba mainly as a “sink”, receiving colonists or immigrants, and Hispaniola acts predominantly a source of colonists. This highlights the importance of water currents in determining the prevailing dispersal direction and warrants future explorations of the impacts of modeling water currents in Caribbean historical biogeography.

### 4.2 *In situ* diversification, inter-island dispersal, and plausibility of the jump dispersal model

Our findings indicate that *in situ* diversification has played an essential role in shaping high levels of endemism of Caribbean *Exophthalmus* species. *In situ* diversification comprised ~73.4% of the 64 phylogeny-internal biogeographic events, whereas FEJD accounts for the remaining events (~25%, 16/64). *In situ* diversification is recognized as an important driver of species diversity and has been documented for many island lineages (e.g., Bennett & O’Grady, 2013; Bruyn et al., 2014; Cowie, 1995; Matos-Maraví et al., 2014). High rates of *in situ* diversification on islands may be due to several factors such as an abundance of open niches on islands (Gillespie et al., 2012), within-island vicariance (Matos-Maraví et al., 2014), environmental and topographic variability (Paulay, 1994), large area size (Losos & Schluter, 2000), and long emergent history (Bruyn et al., 2014). Several Greater Antillean entimine genera such as *Apotomoderes, Apodrosus*, and *Lachnopus* show distinct biogeographic patterns such as high-level of endemism and apparent within-island diversification that merit further attention (e.g., Franz, 2010; Girón & Franz, 2010, 2012).

How plausible is the dispersal of *Exophthalmus* species across the oceanic barriers? Several studies of diverse lineages show that rafting on flotsam is a possible means of longdistance oceanic over-water dispersal (Cheng & Birch, 1978; Donlan & Nelson, 2003; Heatwole & Levins, 1972). In particular, Heatwole & Levins (1972) found pieces of flotsam with live adult weevils in the waters around the islands of the Puerto Rican bank. Franz et al. (2009) invoked dispersal to explain the distribution of widespread weevil species on Mona Island. Over-water oceanic dispersal is also a likely colonization mechanism for weevils of the Galapagos Islands (Peck, 2001; Sequeira et al., 2000), and of the South Indian Ocean Province Islands in the sub-Antarctic region (Chown, 1994). Many entimine lineages are known to have become established in areas outside their native ranges (e.g., O’Brien et al., 2006; Pinski et al., 2005; Woodruff, 1985), providing evidence that colonization of new habitats is possible. Additionally, species of the EGC appear to be generalist herbivores (Dixon, 1954), a habit may further promote their capacity for dispersal and colonization.

### 4.3 Island-to-continent colonization

A surprising outcome of our study is the colonization of Central American mainland from the Caribbean (Fig. 3). The most fitting biogeographic estimate (DIVALIKE+J on M1 model constraint) infers a jump dispersal from Hispaniola to Central America at 18.1 Ma (Fig. 3). Until recently it had been held that colonization of islands from continents is unidirectional, but evidence supporting invasion of continents by island species is emerging (reviewed in Bellemain & Ricklefs, 2008). Islands are frequently thought to be easier to invade because island communities have a lower diversity, their ecological spaces are less densely packed, and interspecific competition is considered less intense (Bellemain & Ricklefs, 2008; Nicholson et al., 2005). Conversely, continents harbor more species and hence more potential colonists. Furthermore, continental populations are selected for increased competitive ability, making colonists more likely to become established on islands. The reverse colonization from the Caribbean to Central America in the our focal taxon has produced widespread, species-rich continental communities, suggesting that island species are not fundamentally disadvantaged as colonizing competitors in comparison their continental counterparts.

### 4.4. Implications on Caribbean biogeography

Our analyses add to the still limited number of model-based inferences focusing on the historical biogeography of Caribbean invertebrates (see Lewis et al., 2015; Matos-Maraví et al., 2014; McHugh et al., 2014; Oneal et al., 2010). In that greater context of understanding diversification in the Caribbean archipelago, our results convey new insights which we discuss in the following.

We may recognize three major Caribbean biogeography hypotheses with relevance to our analysis. (1) The vicariance hypothesis specifies that Caribbean biota originated on a proto-Antillean arc once connected to Neotropical continents in the late Cretaceous (95–65 Ma), which subsequently drifted eastwards to its current position (Rosen, 1975, 1985). (2) The dispersal hypothesis proposes that organisms colonized Caribbean islands mainly from South America via oceanic over-water dispersal throughout the Cenozoic (65.5 Ma to present) and continued dispersal between islands (e.g., Hedges et al., 1992; Heinicke et al., 2007; reviewed in Hedges, 2001 & 2006). (3) The “GAARLandia” hypothesis maintains that the Greater Antilles and northern South America were briefly connected by a Greater Antilles + Aves Ridge land span during the Eocene–Oligocene transition (35–33 Ma) (Iturralde-Vinent & MacPhee, 1999).

Perhaps surprisingly, the over-water dispersal hypothesis best aligns with our preferred biogeographic model reconstruction (i.e., DIVALIKE+J with the M1 model constraint). Both the vicariance and the GAARLandia hypotheses are more difficult to reconcile with this reconstruction due to incongruences in the timing of diversification. Moreover, our inferred scenario of Antillean species colonizing Central America, or of a paraphyletic Caribbean grade, cannot be reconciled with the ancient vicariance hypothesis which predicts sister-group relationships between Central America and Caribbean lineages. Evidence of repeated inter-island jump dispersal events in the *Exophthalmus*, occurring from 25-10 Ma, poorly aligns with an auxiliary postulate of the GAARLandia hypothesis, i.e., island-to-island vicariance (Iturralde-Vinent & MacPhee, 1999, p. 52). Indeed dispersal over oceanic water gaps appears to be a commonly inferred biogeographic process in the recent literature studying various island systems (Balke et al., 2009; Bell et al., 2014; Clayton et al., 2009; Schweizer et al., 2010).

Could our results convincingly reject the ancient vicariance and the GAARlandia hypotheses? We caution against such interpretations. Compared to the over-water dispersal hypothesis, these two hypotheses have more specific predictions that might be best applied to certain lineages whose ages better fit to the time windows stated by these hypotheses. Other issues such as uncertainty and errors in molecular dating could also alter our understanding of the timing of divergences of the pertinent taxa. Unaccounted South American species will need to be studied in the future to produce a complete picture of the biogeographic history of the EGC. We also echo Hedges (2001) that these hypotheses are not necessarily mutually exclusive of one another. The intricacies of the predictions of the hypotheses need to be teased out in relation to the biogeographic history of the taxa being studied.

## 5. CONCLUSION

Our study sheds new light on the historical biogeography and evolution of a group of weevils with a large radiation in the Caribbean archipelago and Neotropical mainland. Given the strong effects of including FEJD and other model parameters, our analysis of highlights a great need for comprehensive model exploration and statistically mediated data-to-model fit optimization in support of reliable historical biogeographic inferences. Among other surprising insights related to the timing and mode of evolutionary radiation for this lineage, we have uncovered an island-to-continent colonization event, thereby providing further evidence that islands can be an important source of continental diversity. We furthermore show that the endemic diversity of the focal lineages in the Caribbean region is largely the outcome of *in situ* diversification. Many weevil lineages have diversified extensively on islands or island-like habitats (Girón & Franz, 2012; Paulay, 1985; Riedel et al., 2014; Samuelson, 2003; Sequeira et al., 2008; Setliff, 2007; Tänzler et al. 2014, Toussaint et al., 2015; Van Dam & Matzke, 2016), and thus are highly suitable for future research on island evolution and biogeography.

## Abbreviation

EGC, *Exophthalmus* genus complex; FEJD, founder event jump dispersal; Ma, million-years ago; Myr, million years

## ACKNOWLEDGEMENTS

We thank Robert Anderson (Canadian Museum of Nature) for contributed specimens and Charles O’Brien (Green Valley, Arizona) for assistance with species identifications. Anyi Mazo-Vargas and Jennifer Girón assisted with early field work and the former also contributed a portion of the molecular data. Andrew Johnston and Salvatore Anzaldo helped with obtaining images of reference collections. Steven Davis (University of Kansas) and Conrad Labandeira (Smithsonian National Museum of Natural History) arranged fossil loans. Will Sides, Lin Pan, and Kevin Wang (Arizona State University; all SOLUR students) photographed specimen. Research support granted by various funding agencies is kindly acknowledged, as follows: National Science Foundation [grants DEB–1155984 (NMF), EF– 1207371 (NMF), EF-0832858 (NJM), and DBI-1300426 (NJM)]; U.S. Department of Agriculture Agreement [No. 58-1275-1-335 (NMF)]; and U.S. Department of Homeland Security and The University of Tennessee, Knoxville (NJM).

## SUPPLEMENTARY INFORMATION

Model selection in statistical historical biogeography of Neotropical insects—the *Exophthalmus* genus complex (Curculionidae: Entiminae).

Guanyang Zhang, Usmaan Basharat, Nicholas Matzke, Nico M. Franz

**Supplementary method descriptions**

<u>Phylogenetic placements of South American species</u>

12 species of the *Exophthalmus* genus complex are previously recorded from South America, accounting for ~9% of the described diversity of the *Exophthalmus* genus complex. Seven species, namly, *Exophthalmus consobrinus* (Marshall, 1922), *E. annulonotatus* (Waterhouse, 1879), *E. crassicornis* Kirsch, 1868 and *E. parentheticus* (Marshall, 1922), *E. jekelinaus* (White, 1858), *E. sulcicrus* Champion, 1911 and *E. agrestis* can be placed into a predominantly Central American clade (Figure 1 in Franz 2012 and Figure 1 in the current study). These species possess a synapomorphic character that unites the Central American clade, i.e., dorso-lateral fovea anterior to eyes. Four of these species, *E. consobrinus, E. annulonotatus, E. crassicornis* and *E. parentheticus* share uniquely patterned yellow stripes, which are arranged as a hexagon or symmetrical brackets, formed by scales on elytra. These species could be united with one of the Panamanian species (labeled as “*Exophthalmus* aff. annulonotatus sp.GZ73”) in the molecular phylogeny that closely resembles *E. annulonotatus*¸ which also has the uniquely patterned stripes on elytra. Four species are recorded from both Central and South Americas: *E. crassicornis, E. sulcicrus, E. jekelianus* and *E. agrestis*. In the current phylogeny we did not have specimens of these species. There is one species from Panama that we tentatively identified to be *E. jekelianus*. Our ongoing research indicates that *E. jekelianus* may represent a species complex and it is not clear if specimens from Central and South Americas previously determined to belong to this species actually represent the same species. We thus refrained from coding this species as widespread in both Central and South America. With regard to the remaining five species previously recorded from South America, we do not have museum or fresh specimens for four species: *E. humilis* (Gyllenhal, 1834), *E. pulchellus* Boheman, 1834, *E. murinus* (Rosenschoeld, 1840) and *Tropirhinus novemdecimpunctatus* (Fabricius, 1775) and their descriptions do not contain adequate information to inform their phylogenetic placement. Another species *E. cinerascens* (Fabricius, 1792) was reported from both Hispaniola and French Guiana. This species was originally described from Hispaniola. A specimen of this species from Hispaniola was included in our study and we are confident about its identification. The publication (Schoenherr et al., 1834) that reported the occurrence of *E. cinerascens* from French Guiana did not provide any details regarding its identification and since that time there has not been any record of this species from French Guiana. We thus consider this occurrence record dubious and the geographic coding of *E. cinerascens* was limited to Hispaniola.

<u>DNA data partition and model selection using PartitionFinder (version 1.1.1)</u>

Three partitions were identified by running an analysis with ParitionFinder. The first partition corresponds to the 12s gene fragment. The second partition includes the 28s gene fragment, all codon positions of arginine kinase (AK), the first and second codon positions of COI and COII, and all codon positions of EF 1-alpha. The third partition pertains to the third codon positions of COI and COII. The partitions scheme is further elaborated in the following, with detailed annotations of the genes and codon positions.

Settings used in PartitionFinder

alignment: ./Entiminae_91T_phylip.phy

branchlengths: unlinked

models: GTR+I, GTR+G, TrNef+G, TrN+G, TrN+I, TrNef+I, HKY, K80, HKY+I+G, K80+I, SYM+G, SYM+I, K80+G, TrNef, GTR, TrNef+I+G, HKY+G, SYM+I+G, TrN, HKY+I, SYM, GTR+I+G, K80+I+G, TrN+I+G

model_selection: bic

search: greedy

Best partitioning scheme

Scheme Name: step_11

Scheme lnL: −69322.50388

Scheme BIC: 143437.064463

Number of params: 566

Number of sites: 4753

Number of subsets: 3

Subset | Best Model | Subset Partitions | Subset Sites | Alignment

1. | GTR+I+G | Gene1 12s | 1-588 | ./analysis/phylofiles/c8dbd8c9e371bd88bf8cc34c53a08063.phy
2. | GTR+I+G | Gene2 28s, Gene3 AK pos1, Gene3 AK pos2, Gene3 AK pos3, Gene4 COII pos1, Gene4 COII pos2, Gene5 COI pos1, Gene5 COI pos2, Gene6 Ef1a pos1, Gene6 Ef1a pos2, Gene6 Ef1a pos3 | 589-1429, 1430-2176\3, 1431-2176\3, 1432-2176\3, 2177-2904\3, 2178-2904\3, 2905-4205\3, 2906-4205\3, 4206-4753\3, 4207-4753\3, 4208-4753\3 | ./analysis/phylofiles/4c5f5027004d6ee804685f2502ec556c.phy
3. | GTR+G | Gene4_COII_pos3, Gene5_COI_pos3 | 2179-2904\3, 2907-4205\3 | ./analysis/phylofiles/5783202a193c474a5f004a983d52c370.phy

Scheme Description in PartitionFinder format

Scheme_step_11 = (Gene1 12s) (Gene2 28s, Gene3 AK pos1, Gene3 AK pos2,

Gene3 AK pos3, Gene4 COII pos1, Gene4 COII pos2, Gene5 COI pos1, Gene5 COI pos2,

Gene6 Ef1a pos1, Gene6_Ef1a_pos2, Gene6_Ef1a_pos3) (Gene4_COII_pos3,

Gene5_COI_pos3);

Partition definitions

DNA, p1 = 1-588;

DNA, p2 = 589-1429 1430-2176\3 1431-2176\3 1432-2176\3 2177-2902\3 2178-2902\3 2903-4201\3 2904-4201\3 4202-4747\3 4203-4747\3 4204-4747\3;

DNA, p3 = 2179-2902\3 2905-4201\3;

<u>Fossil calibrations and age prior parameters</u>

We followed the best practices recommended by Parham et al. (2012) to perform fossil calibrations. We determined the phylogenetic placements of fossils based on apomorphic characters proposed in Franz (2012). Fossil specimen with accession number “NMNH/Woodruff #9768 (National Museum of Natural History, USA)” was used to calibrate the clade “*Compsoricus maricao–Tropirhinus* sp.GZ50”. This fossil possesses at least two synapomorphies of this clade: (1) elytral apices acutely projecting (character 67, state 1 in Franz, 2012); and (2) rostrum in lateral profile slightly arched and tumescent in mid region of dorsal surface (character 9-1 in Franz 2012). Fossil specimen with accession number “NMNH, Woodruff #9774” was used to calibrate the clade “*Exophthalmus roseipes–Diaprepes marginicollis”*. This fossil specimen exhibits three synapomorphic conditions of this clade: (1) a tricarinate rostrum, where the lateral angles of rostrum are distinctly angulate with slight elevation (character 17-1), (2) scrobe passing over eye (character 23-0); and (3) pronotum in dorsal profile with small, shallow, densely arranged, irregularly shaped and spaced concavities (character 45-1). The last specimen (KU-NHM-ENT DR-888, The University of Kansas) was determined to be part the clade “*Scelianoma elydimorpha*–*Artipus monae*” as shown in Figure 2 in Franz (2012) based on these characters: (1) hindwings absence (humeri greatly reduced) (character 83-1); (2) scrobe passing over eye (character 23-0); and (3) occipital sutures anteriorly extending to subapex of rostrum, anteriorly ascending and visible in lateral profile (character 26-1). In the current analysis, *S. elydimorpha* and *A. monae* did not form a clade. According to the reconciliation method proposed by
Parham et al. (2012) (see Figure 2 therein), this fossil can be used to calibrate nodes leading to either of these species or the most recent common ancestor of a clade including both species. As there was no further evidence to base on to determine which of these two species the fossil should be placed with for calibration, we opted to go with the second option. And the clade containing both *S. elydimorpha* and *A. monae* is the “Entiminae sp.GZ34– Entiminae sp. 6” clade. All three amber fossils are from Dominican Republic and were determined to be from the Early Miocene (Burdigalian, 16-20 Ma) by experts working on these fossils (Steve Davis and David Grimaldi, American Museum of Natural History; unpublished data). The prior parameters of the lognormal distributions of the ages are specified as follows. The offset, mean and standard deviation of the lognormal distributions were set so that the 95% interval (2.5% to 97.5% quantiles) corresponded to the minimum ages of the fossils and soft maximum bounds that are reasonably larger than the possible maximum age of the fossils. The calibration based on the KU fossil (KU-NHM-ENT DR-888) for the “Entiminae sp.GZ34– Entiminae sp. 6” clade was given a lognormal prior (in Myr) of log(Mean) = 1.8, log(Stdev) = 1.5 and offset = 15.7, (2.5–97.5% quantiles = 16.0–130.1). The clade “*Compsoricus maricao– Tropirhinus* sp.GZ50” was set have log(Mean) = 1.8, log(Stdev) = 1.3 and offset = 15.5 (2.5– 97.5% quantiles = 16.0–92.8), based on the NMNH fossil Woodruff #9768. The “*Exophthalmus roseipes*–*Diaprepes marginicollis*” clade was given log(Mean) = 2.82, log(Stdev) = 0.5, and offset = 9.7 (2.5–97.5% quantiles = 16.0–54.4) based on NMNH the fossil Woodruff #9774.

---

**Table S1.**
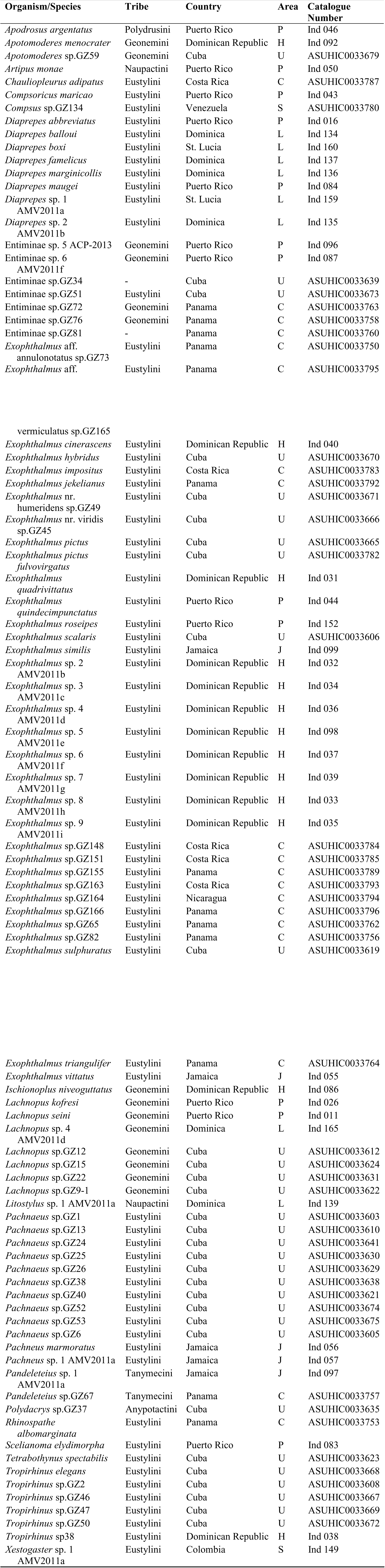
Voucher specimen information. Area code: C-Central America, H-Hispaniola, L-Lesser Antilles, J-Jamaic, P-Puerto Rico, S-South America, U-Cuba. Some specimens were used in a previous study (Mazo-Vargas, unpublished) and were not assigned ASUHIC specimen voucher numbers (given an ‘ Ind’ number instead).

**Table S2.**
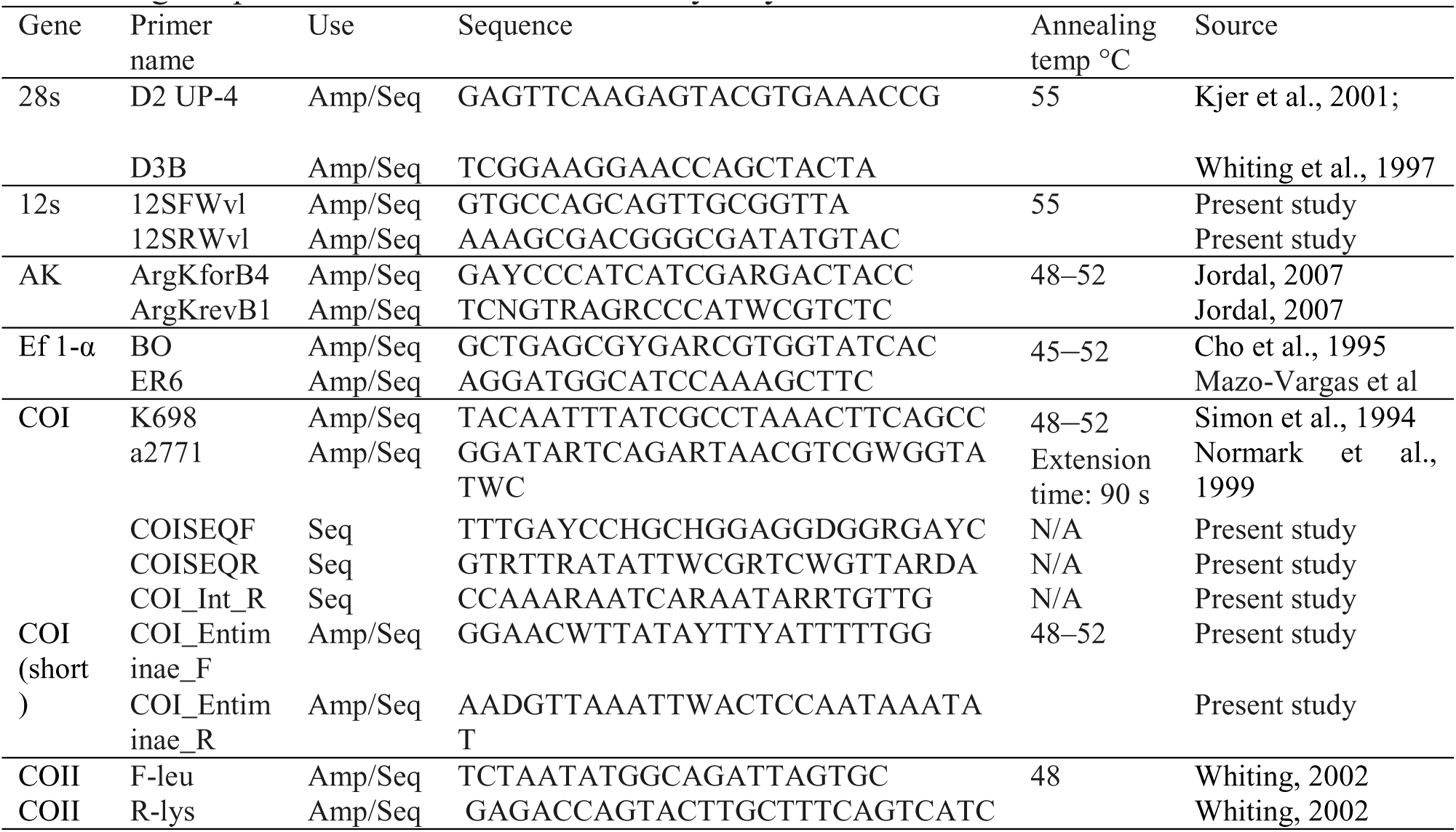
PCR primers and programs. Abbreviations: AK – Arginine kinase Ef 1-α – Elongation Factor 1-alpha COI – Cytochrome c Oxidase subunit I COII – Cytochrome c Oxidase subunit II Amp – amplification Seq – sequencing PCR program follow this general routine: (1) denaturation – 5 min; (2) 35 cycles of: denaturation – 60 s, annealing – 60 s, temp: 48-52 °C, extension – 60 s; and (3) final extension – 7 min. Annealing temperature and extension time may vary and are indicated in the table.

**Table S3.**
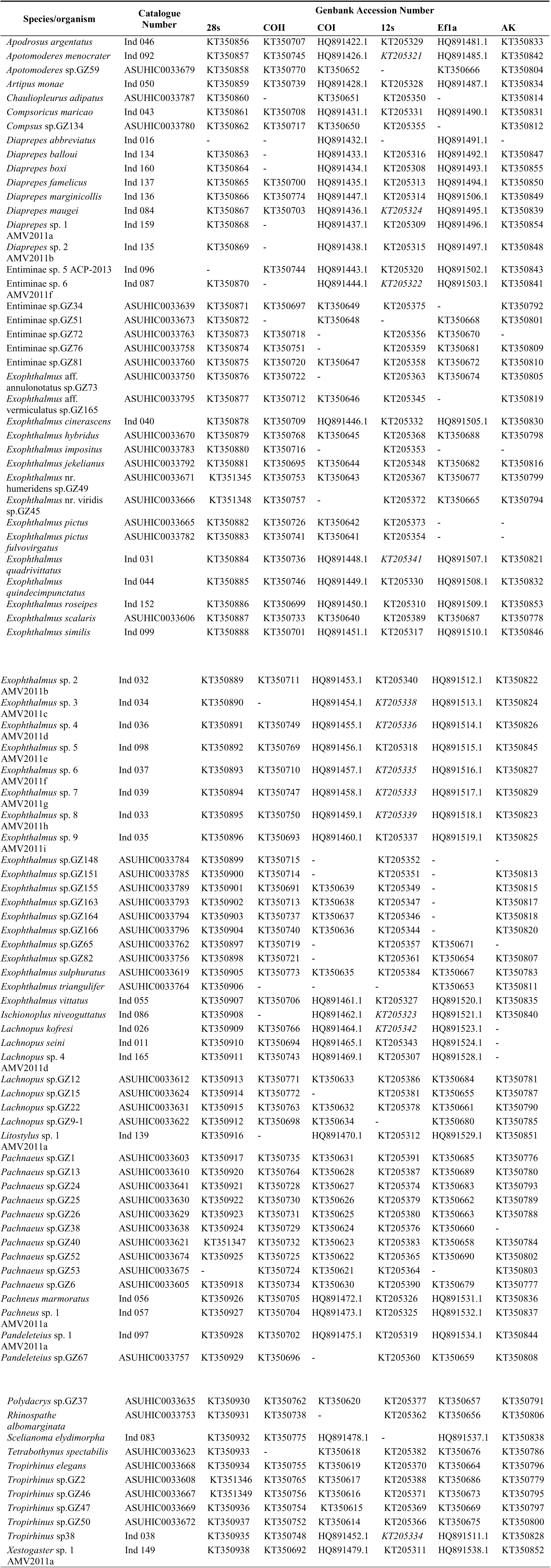
Genbank accession numbers. Abbreviation: AK – Arginine kinase, Ef 1-α – Elongation Factor 1-alpha, COI – Cytochrome c Oxidase subunit I, COII – Cytochrome c Oxidase subunit II

**Table S4.** Summary of dating analyses with different clock and age prior settings.

| # | clock model | Age prior | Median age estimates (95% HPD) |  |  |  |
| --- | --- | --- | --- | --- | --- | --- |
|  |  |  | *Exophthalmus* genus complex | "Entiminae sp.GZ34–Entiminae sp. 6" clade | "*Compsoricus maricao*–*Tropirhinus* sp.GZ50" clade | "*Exophthalmus roseipes*–*Diaprepes marginicollis*" clade |
| 1 | linked | Log normal | 24.57 (18.01-33.38) | 28.03 (19.94-38.47) | 20.27 (15.64-27.49) | 18.01 (13.12-24.11) |
| 2 | linked | Exponential | 25.28 (20.16-36.64) | 29.15 (22.54-42.81) | 21.10 (16.03-30.34) | 18.30 (16.00-26.11) |
| 3 | unlinked | Log normal | 22.93 (17.94-31.98) | 24.45 (18.70-34.22) | 18.85 (15.58-26.05) | 17.79 (13.90-24.56) |
| 4 | unlinked | Exponential | 24.06 (19.48-34.85) | 25.79 (20.38-37.94) | 19.94 (16.00-28.79) | 18.37 (16.00-26.36) |

**Table S5.** Dispersal rate multipliers between areas specified for the M1 constraint.

| Area | S | L | P | H | J | U | C |
| --- | --- | --- | --- | --- | --- | --- | --- |
| Dispersal rate multiplier | 1 | 1 | 1 | 1 | 1 | 1 | 1 |
|  | 0.5 | 1 | 1 | 1 | 1 | 1 | 1 |
|  | 0.5 | 0.5 | 1 | 1 | 1 | 1 | 1 |
|  | 0.5 | 0.5 | 0.5 | 1 | 1 | 1 | 1 |
|  | 0.5 | 0.5 | 0.5 | 0.5 | 1 | 1 | 1 |
|  | 0.5 | 0.5 | 0.5 | 0.5 | 0.5 | 1 | 1 |
|  | 0.5 | 0.5 | 0.5 | 0.5 | 0.5 | 0.5 | 1 |

**Figure S1.**
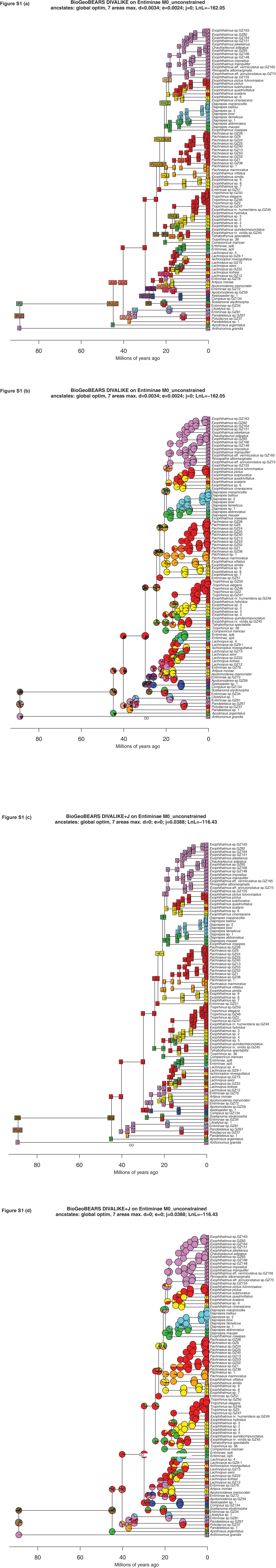
Ancestral range estimations of the *Exophthalmus* genus complex and outgroups using the DIVALIKE model with the M0 model constraint (equal transition between any areas and across time, or essentially not constrained). The dated phylogeny is based on the maximum clade credibility tree generated from a BEAST analysis with the Central American ingroup species constrained to be monophyletic. Abbreviation: C—Central America, H—Hispaniola, J—Jamaica, L— Lesser Antilles, P—Puerto Rico, S—South America, and U—Cuba. **(a)** Single most probable ranges shown, without jump dispersal. **(b)** Relative probabilities of all possible ranges shown in pie charts, without jump dispersal. **(c)** Single most probable ranges shown, with jump dispersal. **(d)** Relative probabilities of all possible ranges shown in pie charts, with jump dispersal.

**Figure S2.**
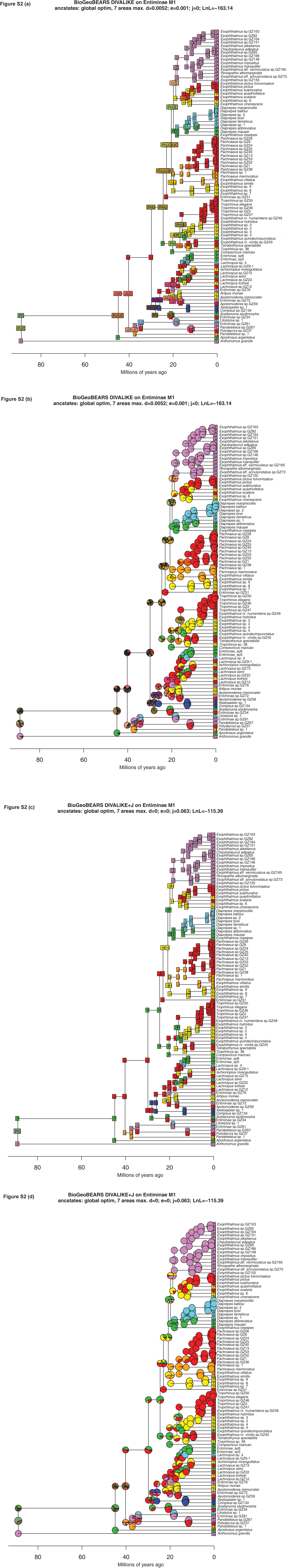
Ancestral range estimations of the *Exophthalmus* genus complex and outgroups using the DIVALIKE model with the M1 model constraint (higher transition rate assigned to dispersals in the same direction as prevailing water currents in the Caribbean). The dated phylogeny is based on the maximum clade credibility tree generated from a BEAST analysis with the Central American ingroup species constrained to be monophyletic. Abbreviation: C—Central America, H—Hispaniola, J—Jamaica, L—Lesser Antilles, P—Puerto Rico, S—South America, and U—Cuba. **(a)** Single most probable ranges shown, without jump dispersal. **(b)** Relative probabilities of all possible ranges shown in pie charts, without jump dispersal. **(c)** Single most probable ranges shown, with jump dispersal. **(d)** Relative probabilities of all possible ranges shown in pie charts, with jump dispersal.

**Figure S3.**
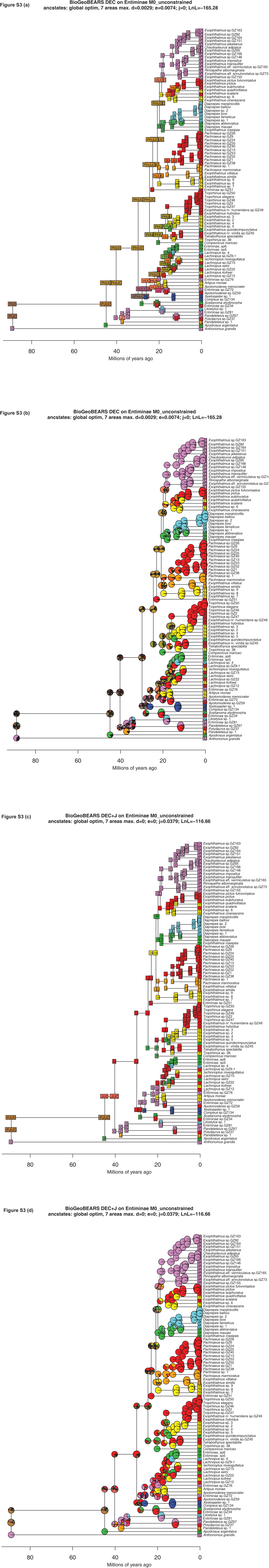
Ancestral range estimations of the *Exophthalmus* genus complex and outgroups using the DEC model with the M0 model constraint (higher transition rate assigned to dispersals in the same direction as prevailing water currents in the Caribbean). The dated phylogeny is based on the maximum clade credibility tree generated from a BEAST analysis with the Central American ingroup species constrained to be monophyletic. Abbreviation: C—Central America, H— Hispaniola, J—Jamaica, L—Lesser Antilles, P—Puerto Rico, S—South America, and U—Cuba. **(a)** Single most probable ranges shown, without jump dispersal. **(b)** Relative probabilities of all possible ranges shown in pie charts, without jump dispersal. **(c)** Single most probable ranges shown, with jump dispersal. **(d)** Relative probabilities of all possible ranges shown in pie charts, with jump dispersal.

**Figure S4.**
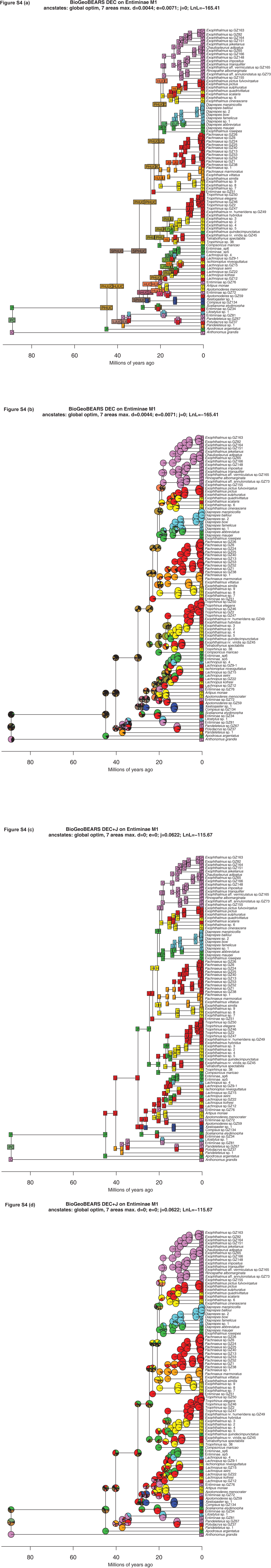
Ancestral range estimations of the *Exophthalmus* genus complex and outgroups using the DEC with the M1 model constraint (higher transition rate assigned to dispersals in the same direction as prevailing water currents in the Caribbean). The dated phylogeny is based on the maximum clade credibility tree generated from a BEAST analysis with the Central American ingroup species constrained to be monophyletic. Abbreviation: C—Central America, H— Hispaniola, J—Jamaica, L—Lesser Antilles, P—Puerto Rico, S—South America, and U—Cuba. **(a)** Single most probable ranges shown, without jump dispersal. **(b)** Relative probabilities possible ranges shown in pie charts, without jump dispersal. **(c)** Single most probable shown, with jump dispersal. **(d)** Relative probabilities of all possible ranges shown in pie with jump dispersal.

**Figure S5.**
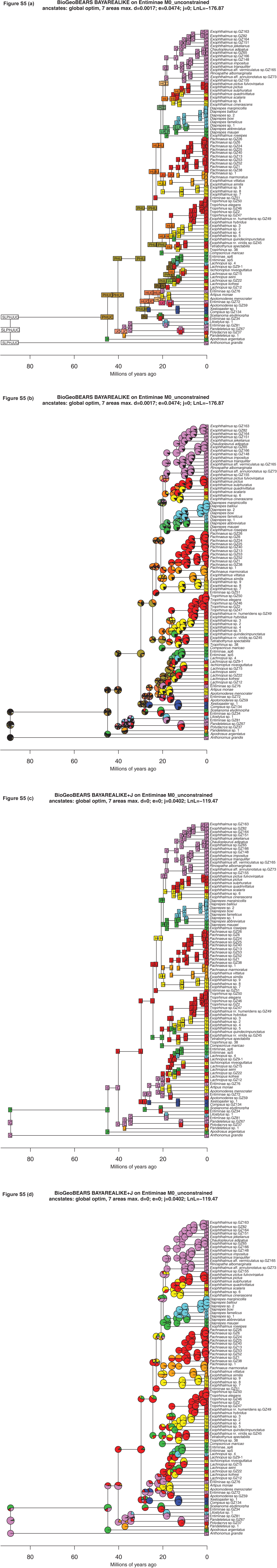
Ancestral range estimations of the *Exophthalmus* genus complex and outgroups the BAYAREALIKE model with the M0 model constraint (higher transition rate assig dispersals in the same direction as prevailing water currents in the Caribbean). The phylogeny is based on the maximum clade credibility tree generated from a BEAST a with the Central American ingroup species constrained to be monophyletic. Abbreviatio Central America, H—Hispaniola, J—Jamaica, L—Lesser Antilles, P—Puerto Rico, S—South A and U—Cuba. **(a)** Single most probable ranges shown, without jump dispersal. **(b)** R probabilities of all possible ranges shown in pie charts, without jump dispersal. **(c)** Single probable ranges shown, with jump dispersal. **(d)** Relative probabilities of all possible shown in pie charts, with jump dispersal.

**Figure S6.**
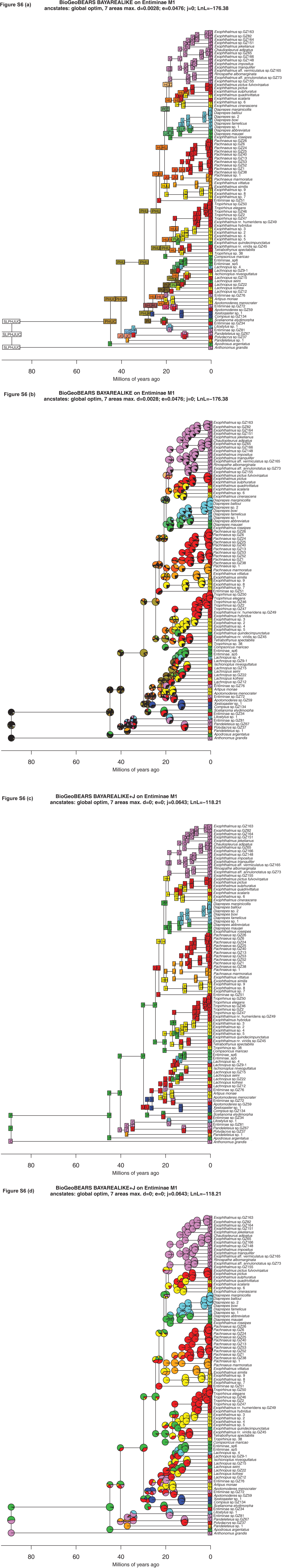
Ancestral range estimations of the *Exophthalmus* genus complex and outgroups the BAYAREALIKE model with the M1 model constraint (higher transition rate assig dispersals in the same direction as prevailing water currents in the Caribbean). The phylogeny is based on the maximum clade credibility tree generated from a BEAST a with the Central American ingroup species constrained to be monophyletic. Abbreviatio Central America, H—Hispaniola, J—Jamaica, L—Lesser Antilles, P—Puerto Rico, S—South A and U—Cuba. **(a)** Single most probable ranges shown, without jump dispersal. **(b)** R probabilities of all possible ranges shown in pie charts, without jump dispersal. **(c)** Single probable ranges shown, with jump dispersal. **(d)** Relative probabilities of all possible shown in pie charts, with jump dispersal.

**Figure S7.**
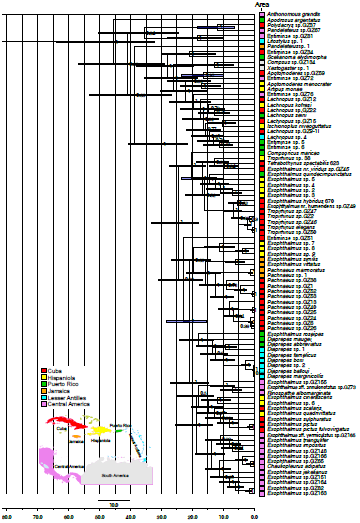
Alternative MCC phylogeny of the *Exophthalmus* genus complex and outgroups where the Central American ingroup species are not constrained to be monophyletic

**Figure S8.**
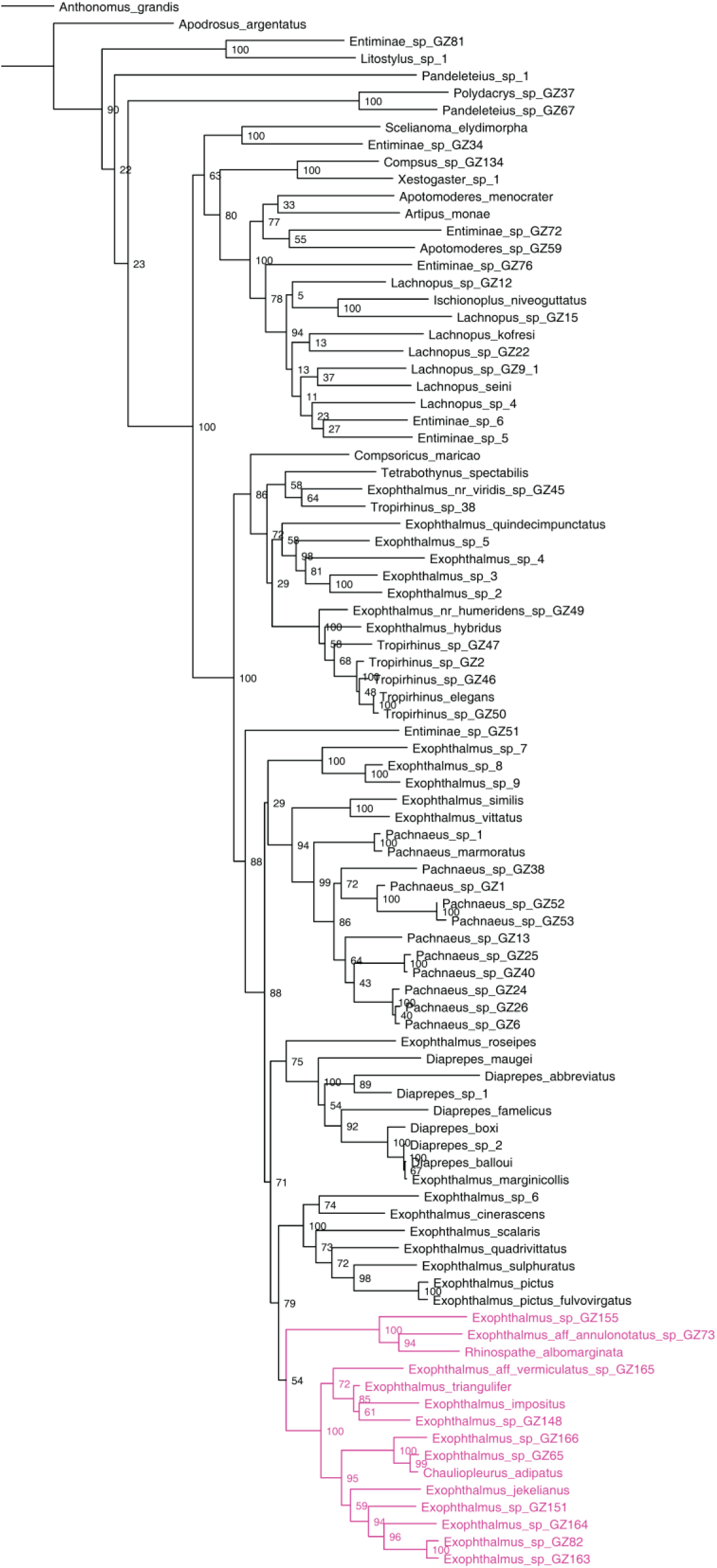
Maximum likelihood phylogeny of the *Exophthalmus* genus complex, which rec the monophyly of Central American species (in magenta).

